# AntiFP2: Genome and Metagenome-Wide Prediction of Antifungal Proteins

**DOI:** 10.64898/2025.12.29.696830

**Authors:** Pratik Balwant Shinde, Shubham Choudhury, Ritu Tomer, Gajendra P. S. Raghava

**Affiliations:** Department of Computational Biology, Indraprastha Institute of Information Technology, Okhla Phase 3, New Delhi-110020, India

**Author notes:** Corresponding Author, Prof. Gajendra P. S. Raghava, Head and Professor, Department of Computational Biology Indraprastha Institute of Information Technology, Delhi Okhla Phase III, New Delhi, India – 110020, Website: http://webs.iiitd.edu.in/raghava/. **Mailing Address of Authors:** Pratik Balwant Shinde (PS), Shubham Choudhury (SC), Ritu Tomer (RT), Gajendra P. S. Raghava (GPSR).

**Keywords:** Antifungal proteins, Genome-wide prediction, Metagenomics, Large language model, Ensemble models, ESM2

## Abstract

The identification of antifungal proteins (AFPs) is crucial for understanding microbial interactions and facilitating the discovery of antifungals on a large scale from genomic and metagenomic data. Most existing computational tools are developed for predicting antifungal peptides rather than proteins. To address this limitation, we developed AntiFP2. A manually curated dataset of experimentally validated AFPs was used to train and develop prediction models using cross-validation techniques. The ensemble strategy, which combined ESM2, BLAST, and MERCI, achieved the best results across independent validation. Beyond prediction, we implemented a complete pipeline for genome- and metagenome-wide AFP screening. AntiFP2 is freely available as a web server, standalone package, and Docker container, enabling scalable and reproducible antifungal protein discovery.

## 1. Background

Fungal pathogens constitute a significant threat to global public health, with invasive mycoses affecting approximately 6.5 million individuals annually and contributing to over 3.8 million deaths worldwide [1]. Fetal fungal infections, despite available treatment, have increased over the past three decades [2]. The increased mortality rate is accompanied by the emergence of multidrug-resistant fungal strains and limitations in existing antifungal drugs, as well as unfavourable pharmacokinetic properties [3].

The study of antifungal proteins (AFPs) has shifted from traditional drug discovery methods to genome and metagenome-based analyses of microbial communities in order to identify novel candidates [4]. Antifungal peptides have been explored as drug candidates for many years, but antifungal proteins have broader biological and ecological significance [5]. Many microorganisms are known to naturally produce proteins that help them survive in environments by suppressing the growth of fungal species [6]. Such microbes may be useful in agriculture by protecting crops against fungal invasion, as well as probiotic candidates in humans. Studying the distribution and diversity of AFPs in genomes and metagenomes enables us to understand the antifungal potential within microbial ecosystems, an area that has not been widely explored in microbiome research [4,7].

However, predicting antifungal proteins from genomic or metagenomic data remains a major challenge. Experimentally screening every protein in the microbial genome is not realistic, and existing computational tools are majorly designed for antifungal peptide identification. [5]. These methods often employ frameshift methods to identify antifungal peptides within proteins using sequence similarity or simple compositional features, which limits their ability to detect novel antifungal proteins, especially in metagenomic datasets where the use of frameshift methods is computationally expensive [8]. To address this gap, there is a need for computational tools that can perform large-scale identification of antifungal proteins on large datasets such as genomes and metagenomes. Such approaches can open new possibilities in microbial ecology, host–microbe interactions, environmental microbiology, and probiotic selection, offering a deeper understanding of how antifungal activity is distributed across microbial communities.

Several computational resources have been developed to support antifungal sequence discovery; however, most are designed primarily for peptide-level prediction rather than full-length proteins. Databases such as CAMPR4, APD3, DBASSP and DRAMP curate experimentally validated antimicrobial peptides, including those with antifungal activity, providing useful reference sets for peptide-focused studies [8–11]. A number of machine-learning and deep-learning models such as Antifp, AFP-MFL, iAFPs-Mv-BiTCN, MLAFP-XN, Deep-AFPpred and DeepAFP further extend these resources by predicting antifungal peptides using compositional features, evolutionary profiles or protein-language-model embeddings [5,12–16]. Although valuable for peptide discovery, these predictors are not inherently designed for full-length antifungal proteins. Macrel (Antimicrobial Peptide Recognition and Evaluation) is another notable tool designed for the genome and metagenome-wide prediction of antimicrobial peptides, including antifungal peptides [17]. Tools such as Antifp, Deep-AFPpred, AMPlify and Macrel offer modules that process proteins, but they do so by fragmenting proteins into short peptides using sliding-frame approaches, subsequently classifying each fragment and making peptide-level predictions [5,16–18]. As a result, their performance depends heavily on the presence of short antifungal motifs rather than holistic protein-level characteristics, limiting their suitability for genome- and metagenome-scale AFP identification. Table 1 summarises the major peptide-focused tools currently available and highlights the methodological emphasis of each.

**Table 1.**
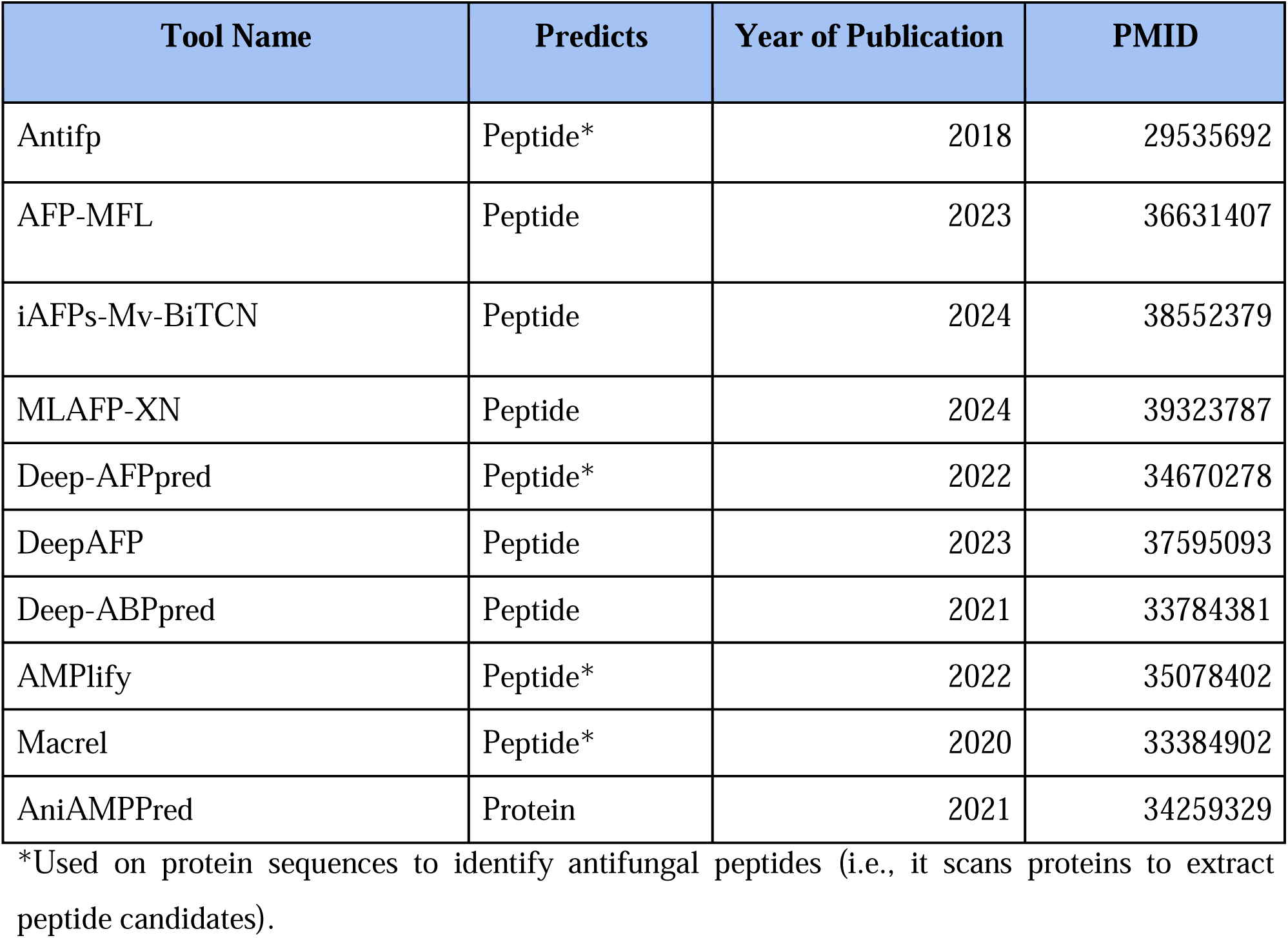
List of existing tools developed for predicting antifungal and antimicrobial peptides and proteins prediction.

| Tool Name | Predicts | Year of Publication | PMID |
| --- | --- | --- | --- |
| Antifp | Peptide* | 2018 | 29535692 |
| AFP-MFL | Peptide | 2023 | 36631407 |
| iAFPs-Mv-BiTCN | Peptide | 2024 | 38552379 |
| MLAFP-XN | Peptide | 2024 | 39323787 |
| Deep-AFPpred | Peptide* | 2022 | 34670278 |
| DeepAFP | Peptide | 2023 | 37595093 |
| Deep-ABPpred | Peptide | 2021 | 33784381 |
| AMPlify | Peptide* | 2022 | 35078402 |
| Macrel | Peptide* | 2020 | 33384902 |
| AniAMPPred | Protein | 2021 | 34259329 |

\*Used on protein sequences to identify antifungal peptides (i.e., it scans proteins to extract peptide candidates).

Current computational resources emphasise on prediction of antifungal peptides, with no dedicated method available for discriminating antifungal and non-antifungal proteins. To address this limitation, we developed a prediction system for proteins spanning 51–535 amino acids. It is constructed on a rigorously curated set of experimentally validated AFPs. Model development and evaluation were conducted in accordance with established standards, including strict redundancy reduction, balanced and imbalanced dataset design, and independent validation.

Furthermore, we implemented an integrated genome- and metagenome-scale prediction architecture that unifies alignment-based modules (BLAST, MERCI) with alignment-free methodologies, including compositional and evolutionary machine learning features, as well as fine-tuned protein language model representations. This multi-tiered framework enables sensitive and robust detection of antifungal proteins across highly diverse sequence and metagenomic datasets.

## 2. Results

In this study, our objective was to develop a prediction model for antifungal protein prediction. Multiple strategies were utilised to complete this task: alignment-based methods, alignment-free methods and ensemble methods. The workflow has been explained diagrammatically in **Figure 2**.

### 2.1 Composition analysis

To characterise sequence-level differences associated with antifungal activity, we compared the amino acid composition (AAC) of antifungal peptides, antifungal proteins, and non-antifungal proteins. The antifungal peptides from the older AntiFP version were also used in this analysis. For each amino acid, the percent composition was calculated. Mann-Whitney U test (p < 0.05) was used to evaluate statistically significant differences between each amino acid composition. This is represented graphically in **Figure 1**, with an error bar indicating 95% confidence intervals. The statistical comparison was performed between antifungal peptides and antifungal proteins, as well as between antifungal proteins and non-antifungal proteins. We observed that cysteine, glycine, and tyrosine have a significantly higher presence in antifungal proteins relative to non-antifungal proteins, whereas glutamic acid, isoleucine, leucine, methionine, serine, and valine were consistently more abundant in non-antifungal proteins. These observations highlight significant biochemical differences associated with antifungal function at the level of amino acids.

**Figure 1.**
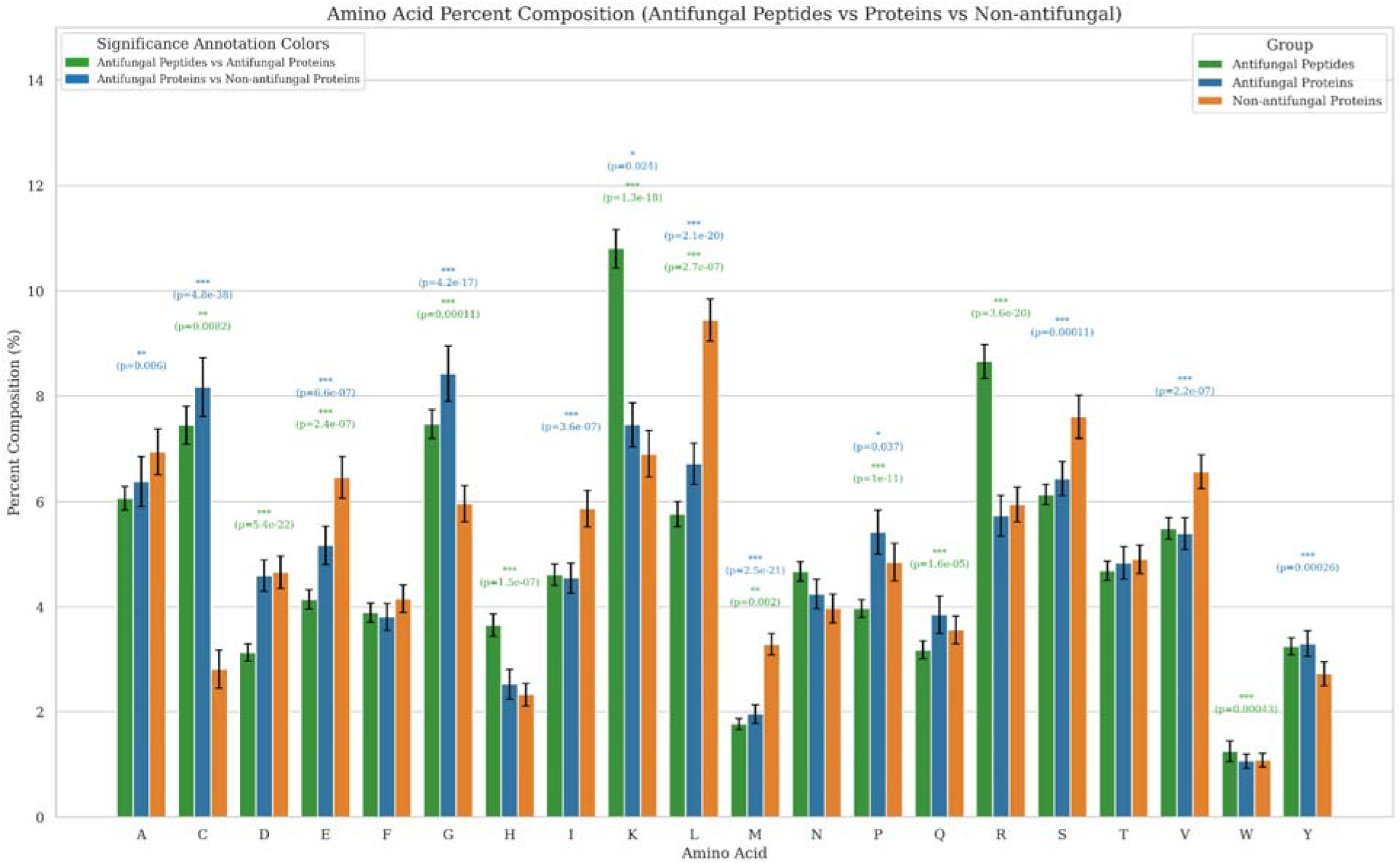
Amino acid composition comparison between the antifungal and non-antifungal datasets, along with statistical significance and 95% confidence interval.

**Figure 2.**
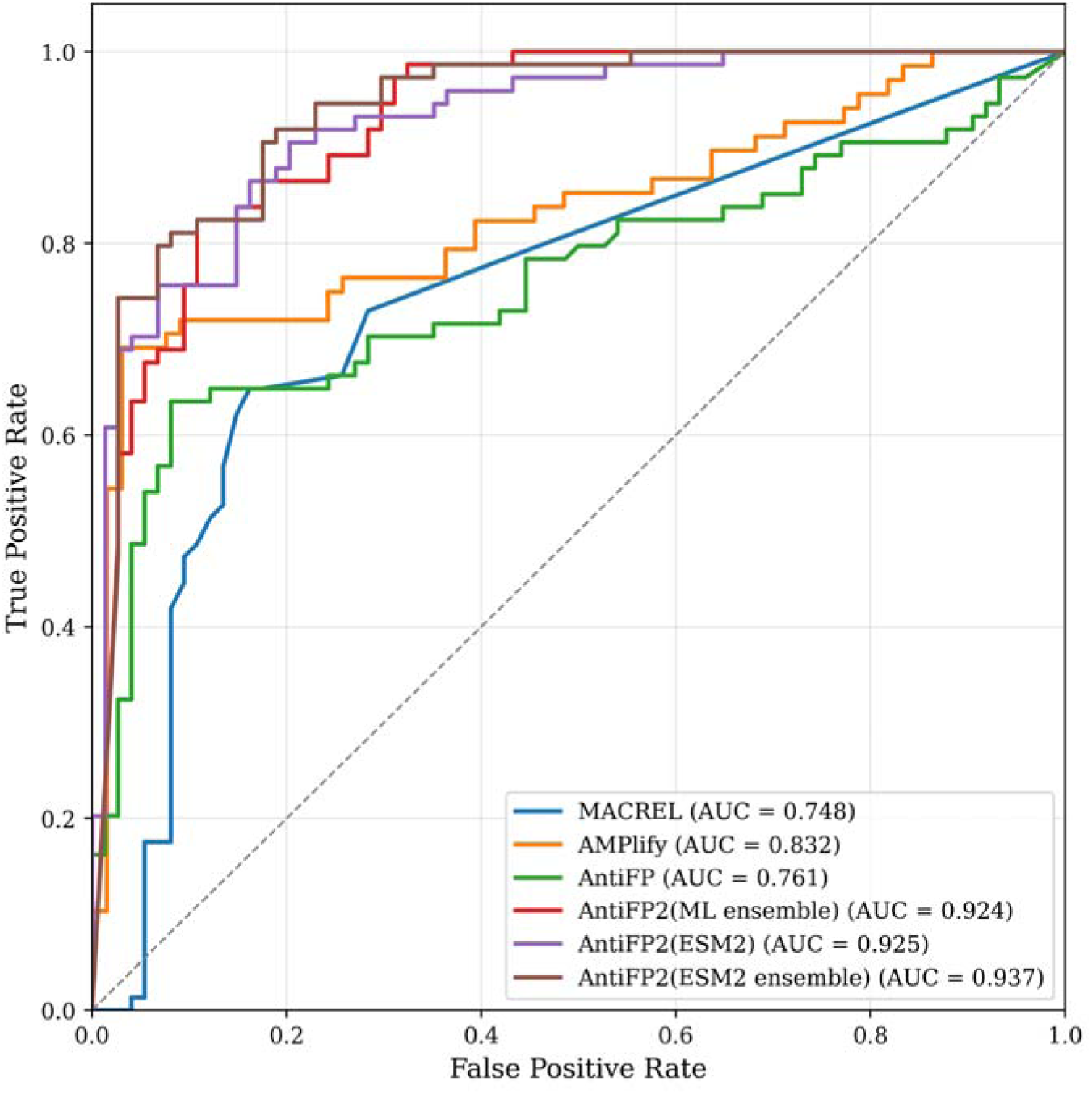
AUROC curve of benchmarking AntiFP2 against other tools.

When comparing antifungal peptides with antifungal proteins, we observed notable differences. Antifungal peptides exhibited a strong enrichment of cysteine, glycine, lysine, and arginine, reflecting the classical cysteine-rich and cationic signatures typical of antimicrobial peptides. Antifungal proteins shared some of these tendencies but with substantially different magnitudes and patterns, indicating that peptide-derived physicochemical biases do not fully translate to full-length proteins. Taken together, these results show that antifungal peptides and antifungal proteins do not share the same sequence pattern. Antifungal proteins have a wider and more variable amino acid distribution, whereas peptides show stronger, more focused patterns driven by specific motifs and positive charge. Because of this difference and because non-antifungal proteins are enriched in several hydrophobic and neutral residues, protein-level antifungal activity cannot be accurately predicted using peptide-based rules or by analysing proteins only through short fragments.

### 2.2 Alignment-Based Methods

Alignment-based methods are widely used in the bioinformatics field for annotating novel proteins based on sequence similarity. We applied two methods: 1. BLAST-based similarity search and 2. Motif-based alignment.

#### 2.2.1. BLAST-based similarity search

BLAST is well known for its fast algorithm for finding sequence similarity between a query sequence and a database, facilitating the annotation of the query sequence. We utilised BLAST to annotate antifungal and non-antifungal proteins, using a local database which was created using our training dataset and the validation dataset was queried against it. The e-value is a measure of the randomness of the match and signifies the quality of alignment. We applied multiple e-values to evaluate the best threshold based on coverage and prediction accuracy. The highest coverage of 99.32% was obtained at E-value 10, but the error rate and accuracy were lowest at 31.97% and 69.03% respectively. E-value was further decreased to achieve a balance between coverage accuracy and error rate. It was observed that BLASTP alone was insufficient to predict all the novel proteins correctly, as the coverage and accuracy had a trade-off. The E-value of 0.01 was selected based on higher accuracy, better coverage and minimal error rate. The detailed results are explained in **Table 2**.

**Table 2.** The performance of BLAST on a balanced dataset at different e-values.

| E-value | Total hits | Correct hits | Incorrect hits | Accuracy % | Coverage % | Error rate % |
| --- | --- | --- | --- | --- | --- | --- |
| 10 | 147 | 100 | 47 | 68.03 | 99.32 | 31.97 |
| 1 | 112 | 83 | 29 | 74.11 | 75.68 | 25.89 |
| 0.1 | 69 | 61 | 8 | 88.41 | 46.62 | 11.59 |
| 0.01 | 57 | 53 | 4 | 92.98 | 38.51 | 7.02 |
| 0.001 | 49 | 45 | 4 | 91.84 | 33.11 | 8.16 |
| 0.0001 | 38 | 35 | 3 | 92.11 | 25.68 | 7.89 |
| 0.00001 | 32 | 29 | 3 | 90.62 | 21.62 | 9.38 |

#### 2.2.2 Motif-based alignment

In motif-based analysis, short conserved sequence patterns or motifs are identified that are characteristic of antifungal proteins. These motifs represent small regions within a protein that may contribute to antifungal activity. To identify such discriminatory patterns, we used the MERCI tool, which detects motifs that occur in one class of sequences (positive dataset) but are absent in the other (negative dataset).

Motif discovery was performed using the antifungal proteins in the training dataset as the positive class. No motifs were extracted from the negative class, as these sequences were selected only on the basis of being “non-antifungal,” without any shared conserved region. We experimented with multiple MERCI parameter combinations, including (i) number of motifs (K), (ii) allowed false-positive rate (FP), and (iii) amino acid classification schemes. The detailed results of all the combinations are available in **Supplementary Table 3**. Increasing the number of extracted motifs, MERCI was unable to identify all antifungal proteins in the validation dataset, indicating that motif signals could only capture a subset of antifungal patterns. Our analysis revealed that specific conserved patterns, including Basic–C–Sulfur–Sulfur, Aliphatic–K–C–Aliphatic, and G–Neutral–C–Aliphatic, were exclusive to antifungal proteins. These unique motifs represent short sequence signatures enriched in proteins with antifungal activity. The detailed table of motifs is available in **Supplementary Table 2.**

Among all tested settings, the best-performing parameter combinations corresponded to K = 8 and K = 9, both achieving an accuracy of 92.86% on the validation set. To avoid overfitting and maintain a concise motif set, we selected the configuration with 8 motifs (K = 8), a false-positive allowance of 10 (FP = 10), and the Koolman–Rohm amino acid classification scheme for subsequent analyses. The eight selected antifungal motifs are listed below:

### 2.3 Alignment-free methods

As per previous observations, the alignment-based methods depend on sequence similarities between the query and database and therefore fail to provide annotations for all the proteins. To overcome these limitations, we employed a range of alignment-free prediction models, primarily utilising artificial intelligence. In this study, we developed machine learning and large language based models.

#### 2.3.1 Machine learning models

To evaluate alignment-free approaches for antifungal protein prediction, we first developed a series of machine learning models using different ML classifiers and feature representations. These included composition-based features generated using Pfeature, evolutionary features derived from PSSM profiles using POSSUM, probability-based encodings, and embedding-based features extracted from large language models (LLMs). A summary of the best-performing models is provided in **Table 3**.

**Table 3.**
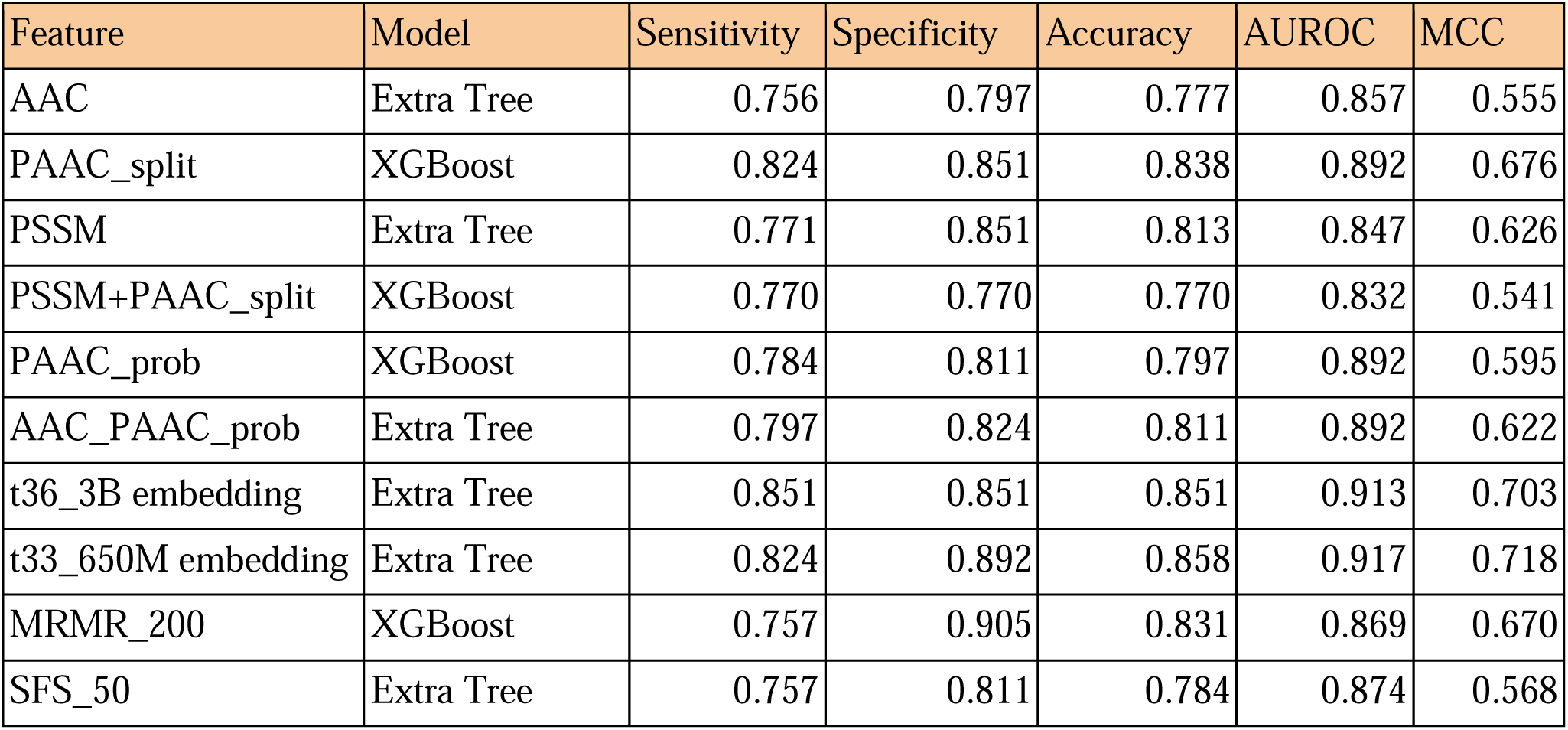
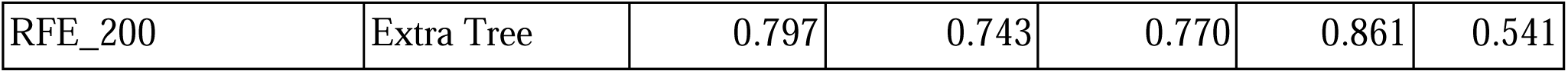
Performance matrix of the best machine learning models trained using the Balanced dataset.

##### 2.3.1.1 Composition-based features

Models trained on standard composition descriptors showed strong baseline performance. The AAC features with Extra Trees classifier achieved highest AUROC of 0.857 and MCC of 0.555, indicating that even simple global composition carries informative signals for antifungal discrimination. To incorporate positional information, each protein sequence was divided into four equal segments, and amino acid composition was computed independently for each segment. This representation improved prediction accuracy considerably with XGBoost, and the PAAC_split features achieved an AUROC of 0.892 and an MCC of 0.676, outperforming the complete sequence models. These results highlight that capturing localised, region-specific composition provides greater discriminatory power than global amino acid frequencies alone. Detailed results of all models are presented in **Supplementary Table 4**.

##### 2.3.1.2 Evolutionary features

Evolutionary information captured using PSSM_COMP features yielded competitive performance, with the Extra Trees classifier achieving a maximum AUROC of 0.847. While useful, these features alone performed slightly below the composition-based feature models. Combining composition-based features with PSSM profiles did not further improve performance; for example, PAAC_split + PSSM achieved an AUROC of 0.832 with XGBoost, which remained lower than that of composition-only models. Detailed results are presented in **Supplementary Table 5.**

##### 2.3.1.3 Probability-based features

Probability-derived feature encodings focus on residue-level distribution patterns rather than absolute composition, providing a complementary perspective to AAC and PAAC_split. Initial experiments using PAAC_split probabilities features resulted in similar performance, with the XGBoost model achieving an AUROC of 0.892 and an MCC of 0.595. To improve predictive strength, we combined the two most informative composition AAC_split and PAAC_split probability features, which did not yield improved performance with the Extra Trees model, with an AUROC of 0.892 and an MCC of 0.622. Detailed results are presented in **Supplementary Table 5.**

##### 2.3.1.4 LLM-based embeddings

To evaluate whether LLM-derived sequence representations improve performance, we extracted fixed embeddings from the ESM2_t33_650M and ESM2_t36_3B models and trained classifiers using these embeddings. Embeddings from t33_650M performed best, achieving an AUROC of 0.917 and MCC of 0.718 using Extra Trees. These embedding-based models outperformed conventional features. The detailed results are provided in **Supplementary Table 3**.

##### 2.3.1.5 Feature selection

Multiple feature selection strategies FSFS, SVC-L1, and MRMR were applied to reduce feature dimensionality and identify key contributors. Although these methods selected informative subsets, feature selection did not substantially improve model performance compared with using the full composition-based feature sets (**Supplementary Table 6**). Most selected features originated from the AAC_split and PAAC_split categories, reinforcing their importance in AFP classification.

#### 2.3.2 Large language-based model

We investigated two complementary approaches for antifungal protein classification using the ESM2 family of large language models. We fine-tuned four variants, t2_12M, t12_35M, t33_650M, and t36_3B, on our curated antifungal training dataset. Performance improved consistently with increasing model size, with the largest model, ESM2_t36_3B, achieving the highest classification MCC and AUROC of 0.637 and 0.925, respectively. These results emphasise the benefit of scaling model capacity for learning complex sequence patterns and support the effectiveness of end-to-end fine-tuning for domain-specific protein classification tasks. The detailed results are provided in **Table 4**.

**Table 4.** Performance of ESM2 fine-tuned models on a balanced dataset.

| Model | Sensitivity | Specificity | Accuracy | AUROC | MCC |
| --- | --- | --- | --- | --- | --- |
| t6_12M | 0.824 | 0.811 | 0.818 | 0.878 | 0.635 |
| t12_35M | 0.824 | 0.811 | 0.818 | 0.898 | 0.635 |
| t33_650M | 0.851 | 0.838 | 0.845 | 0.916 | 0.689 |
| t36_3B | 0.784 | 0.851 | 0.818 | 0.925 | 0.637 |

**Table 5.** Performance of ensemble configurations integrating ML, LLM, and alignment-based methods BLAST(BL) and MERCI (MR).

| Ensemble Configuration | Sensitivity | Specificity | Accuracy | AUROC | MCC |
| --- | --- | --- | --- | --- | --- |
| XGBoost (PAAC_split) | 0.784 | 0.797 | 0.791 | 0.892 | 0.581 |
| XGBoost (PAAC_split) + MR | 0.77 | 0.824 | 0.797 | 0.88 | 0.595 |
| XGBoost (PAAC_split) + BL | 0.784 | 0.797 | 0.791 | 0.899 | 0.581 |
| XGBoost(PAAC_Split) + BL + MR | 0.838 | 0.838 | 0.838 | 0.924 | 0.676 |
| t36_3B | 0.784 | 0.851 | 0.818 | 0.925 | 0.637 |
| t36_3B + BL | 0.797 | 0.851 | 0.824 | 0.932 | 0.650 |
| t36_3B + MR | 0.797 | 0.838 | 0.818 | 0.930 | 0.636 |
| t36_3B + BL + MR | 0.824 | 0.838 | 0.831 | 0.938 | 0.662 |

#### 2.3.3 Ensemble method

In previous studies, it has been observed that ensemble or hybrid methods outperform individual methods in certain scenarios [19,20]. Therefore, to further enhance prediction performance, we implemented an ensemble strategy that integrates alignment-free methods (machine learning models and language models) with alignment-based approaches (BLAST and MERCI). Various ensemble models were constructed by combining ML and LLM models along with alignment-based methods. The detailed performance matrix is provided in the **Supplementary Table 7.**

We first evaluated ensembles built around the XGBoost model trained on PAAC_split features. Adding MERCI motifs to this model decreased performance compared to the baseline (AUROC from 0.892 to 0.880). Replacing MERCI with BLAST improved the AUROC to 0.899, demonstrating that homology information contributes meaningfully to prediction accuracy. The best ML-based ensemble was BLAST, and MERCI were combined with PAAC_split features, achieving an AUROC of 0.924 and an MCC of 0.676.

Ensemble models built using fine-tuned t36_3B and combining BLAST improved the AUROC from 0.925 to 0.932 (MCC from 0.637 to 0.650), while combining t36_3B with MERCI produced an AUROC of 0.930 and MCC of 0.636. The best overall performance was achieved when all three components, t36_3B, BLAST, and MERCI, were combined, resulting in the highest AUROC of 0.938 and MCC of 0.662.

These results indicate that ensemble strategies utilising composition-based features, conserved motifs, and sequence similarity provide more reliable antifungal protein predictions than any single method.

### 2.4 Benchmarking

Benchmarking was performed to compare the performance of our model with that of existing antifungal peptide prediction tools, as no dedicated tool currently exists for identifying antifungal proteins. This evaluation was conducted using a common independent test dataset and standardised performance metrics to ensure a fair and objective comparison.

The older AntiFP tool utilises the frameshift method to predict peptides within a protein; we adopted a unified evaluation strategy, where we calculated the “percentage of antifungal region” by dividing the number of antifungal peptides in the protein by the total number of peptides in the same protein. A protein was considered antifungal if at least 20% of the predicted region was antifungal. The optimal threshold (20%) was chosen by evaluating all cutoffs from 1–100% and selecting the one that minimised the absolute difference between sensitivity and specificity.

The benchmarking results show higher performance of our model compared to existing tools. Macrel achieved high specificity (0.946) but very low sensitivity (0.148), resulting in lower overall performance (AUROC = 0.748, MCC = 0.156). The older AntiFP method, which also relies on frame-shift peptide prediction, performed average (AUROC = 0.761, MCC = 0.418). AMPlify also applies the frame-shift method, which performed better, with a sensitivity of 0.617, specificity of 0.967, and an AUROC of 0.83, achieving an MCC of 0.625. The performance comparison is provided in **Figure 2**.

The fine-tuned ESM2 t36_3B model achieved an AUROC of 0.925 and MCC of 0.637, demonstrating strong and balanced performance. Furthermore, the ESM2-based ensemble model shows the highest AUROC of 0.938, sensitivity of 0.824, and MCC of 0.662. These results confirm the benefits of integrating large language models with similarity-based features for accurate prediction of antifungal proteins. The complete performance matrix is provided in **Supplementary Table 8.**

### 2.5 Realistic and Redundant Dataset Evaluation

To evaluate model robustness under more realistic conditions, we also tested performanc on two additional datasets: a realistic dataset with a 1:10 positive-to-negative ratio, and a redundant dataset consisting of 926 positive proteins prior to CD-HIT filtering and 9260 randomly selected negative proteins, reflecting a highly redundant sequence space. The detailed performance matrix is provided in **Table 7**.

**Table 7.**
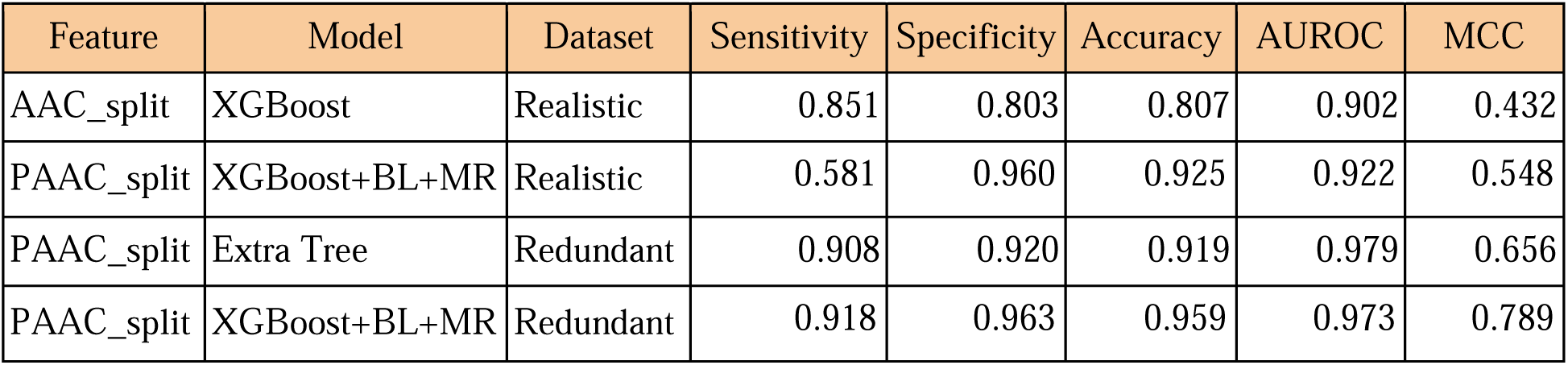
Performance of the models built using a Realistic and Redundant dataset.

In the realistic dataset, the AAC_split model achieved an AUROC of 0.902 and MCC of 0.432. Furthermore, we applied the ensemble model of XGBoost, integrated with BLAST and MERCI, utilising PAAC_split features, which resulted in higher performance. The ensemble achieved an AUROC of 0.922, an MCC of 0.548, and a markedly increased specificity of 0.960, resulting in an overall accuracy of 0.925. Although sensitivity decreased (0.581), the improved specificity and MCC illustrate that hybrid models better handle strong class imbalance by incorporating homology and motif signals.

In the redundant dataset, higher performance was observed in all models, which can be due to the presence of similar sequences in both the training and validation datasets. The Extra Tree model built using PAAC_split features performed better (AUROC of 0.979 and MCC of 0.656) than the ensemble model (XGBoost + BLAST + MERCI) built using the same PAAC_split features (AUROC of 0.973, an MCC of 0.789), but the ensemble model showed higher sensitivity of 0.918 and specificity of 0.963.

### 2.6 Metagenomic Case Study

To assess the practicality of the AntiFP2 metagenome pipeline, we conducted a comparative case study using real-world metagenome samples (SRA: SRR5936131). The sample comprises a total of 27,283,256 reads (27 million), with read lengths ranging from 50 to 101 bp. The Prokka tool predicted 100,170 genes and 70,114 proteins from the sample. The sample was processed using the ESM2 ensemble model pipeline and the ML ensemble pipeline. The ESM2 ensemble pipeline identified 4,253 antifungal proteins in 195,840 seconds (∼54 hours) of runtime. The ML ensemble pipeline demonstrated higher efficiency, identifying 11,982 antifungal proteins in 7,358 seconds (∼2 hours).

Further, we combined the predictions from both pipelines, selected the common hits, and ranked them based on prediction probabilities. The top 5% of these common predictions were designated as high-confidence antifungal protein candidates, totalling 970 proteins. This approach was utilised to create a balance between coverage and precision in the predictions of two independent models. The ML ensemble pipeline offers rapid screening and broader sensitivity, the ESM2 ensemble pipeline provides higher precision. The combination of these models provides a robust and scalable method for antifungal protein prediction from highly complex metagenome samples.

### 2.7 Service to Scientific Community

To provide a user-friendly application we provide sequence-level and genome/metagenome-level AntiFP2 modules. We developed a web server (https://webs.iiitd.edu.in/raghava/antifp2/) that enables users to predict antifungal proteins given a protein sequence, utilising an ML ensemble model. The interface is easy to use, responsive, and device-independent, allowing seamless use across desktops, tablets, and smartphones. A Python library (https://pypi.org/project/antifp2/) has been created, allowing users to run predictions using the ESM2 model, the ESM2 ensemble model, and the ML ensemble model.

For metagenome and genome-level applications, we also developed a fully automated, single-command pipeline that predicts AFP directly from the contigs using all three models. This pipeline is provided as a standalone workflow on GitHub (https://github.com/raghavagps/antifp2) along with a detailed installation guide and step-by-step user instructions. The pipeline is also provided in an environment-independent Docker container (https://hub.docker.com/r/pratik0297/antifp2) with an easy installation option for all users.

## 3. Discussion

Identifying novel antifungal proteins remains a challenge due to the limited availability of experimentally validated sequences across different taxa. In this study, we systematically evaluated alignment-based, alignment-free, and hybrid ensemble strategies to improve antifungal protein prediction.

Alignment-based methods, such as BLAST and MERCI, play a major part in annotation due to their interpretability and precision in detecting homologous or conserved regions within sequences. Therefore, similar to earlier publications, the alignment-based methods have a clear trade-off between coverage and accuracy [19,20]. BLAST achieved high coverage of 99.34% at a relaxed E-value of 10 but with low accuracy (69.03%), while a stringent threshold (E-value ≤ 0.01) improved accuracy to 92.98% but reduced coverage to 38.51%. Similarly, MERCI identified the conserved motifs in the antifungal proteins, but the coverage was very poor. These results emphasise the limitations of alignment-based strategies for detecting novel antifungal proteins.

Alignment-free methods demonstrated greater potential in predicting novel antifungal proteins. Machine learning models built using PAAC_split and probability-based features achieved high prediction performance (both AUROC 0.892). Large language models enhanced the performance even further; the embeddings from the pre-trained ESM2 t33_650M model improved AUROC to 0.917, confirming the ability of the embedding to capture the complex pattern of antifungal sequences. The fine-tuned ESM2 models independently achieved the highest AUROC of 0.925, signifying the importance of domain-specific fine-tuning.

The most effective method was observed to be ensemble models, which integrate alignment-free and alignment-based methods. The ML ensemble achieved an AUROC of 0.924 and an MCC of 0.676, comparable to the ESM2 ensemble model. The ESM2 ensemble model achieved an AUROC of 0.938 and an MCC of 0.662. This emphasises the effect of complementary features being used together to improve the accuracy, interpretability of antifungal protein prediction at a larger scale.

## 4. Conclusion

We provide a robust antifungal protein prediction web server and pipeline for sequence-level and metagenome/genome-level annotation, facilitating the scientific community. The provided modules are designed with an easy-to-use interface and commands, making them accessible to non-computational experts as well. In contrast to existing tools, this method is length-independent and was developed using proteins, making it a unique approach. Further, using the ensemble approach makes it more interpretable and robust for identifying novel antifungal proteins.

## 5. Materials and Methods

### 5.1 Datasets

A reliable prediction model requires a high-quality training dataset; therefore, we included only those antifungal proteins that had prior experimental validation. Antifungal protein sequences were manually curated from several well-established repositories that includes DBAASP [10], CAMP [11], DRAMP [21], APD3 [22], PlantPepDB [23] and UniProt [24]; on 20th August 2024. We considered sequences to be proteins that had an amino acid length of 50 amino acids or above. Based on these filtering criteria, DBAASP had 145 proteins, CAMP had 114 proteins, DRAMP had 377 proteins, APD3 had 127 proteins, and PlantPepDB had 93 proteins. We used a special keyword-based search to retrieve ‘reviewed’ antifungal proteins from UniProt, specifically using the keywords KW-0211 (defensins) and KW-0295 (fungicide), which returned 420 proteins and 1110 proteins, respectively [25]. We also used the term “antifungal” to search the UniProt database, which further returned 1159 proteins. Further, all UniProt protein entries were manually curated to ensure experimental validation and remove sequence similarity-based annotations, and duplicates were removed, resulting in 355 proteins from the UniProt database. We further selected the sequence having a length between 50 to 550 aa, depending on the overall length distribution in the positive dataset. Finally, we obtained 926 unique antifungal proteins called the positive dataset.

We retrieved non-antifungal proteins, referred to as a negative dataset, from the UniProt database for building our classification models. The query used was NOT antifungal, NOT antimicrobial, NOT antibiotic, NOT antimycotic, NOT fungicidal, and NOT fungistatic, resulting in 561,345 ‘reviewed’ proteins as of August 20, 2024. After applying the length filter and removing duplicates, 4,61,952 proteins remained. To remove highly similar sequences, we applied CD-HIT [26] with a 40% sequence identity threshold to both the antifungal and non-antifungal datasets.

After removing the redundancy, the dataset contains 368 antifungal proteins (positive) and 69,467 non-antifungal proteins (negative). Based on this analysis, we created three datasets with different ratios of positive and negative proteins, as described in Figure 3.

- **Balanced dataset:** This dataset consists of 368 positive proteins and 368 negative proteins, randomly selected from the 69,467 non-redundant negative sequences obtained after CD-HIT filtering. Most prediction models in this study were developed using this dataset, unless specified otherwise.
- **Realistic dataset:** This dataset contains the same 368 positive proteins as in the balanced dataset, along with 3,680 negative proteins (1:10 ratio). The negative proteins were randomly selected from the non-redundant negative pool to simulate a more realistic class imbalance.
- **Redundant dataset:** This dataset includes all 926 experimentally validated antifungal proteins prior to redundancy removal. In addition, 9,260 non-antifungal proteins were randomly selected from the non-redundant negative pool. This dataset reflects scenarios where sequence redundancy is retained.

**Figure 3:**
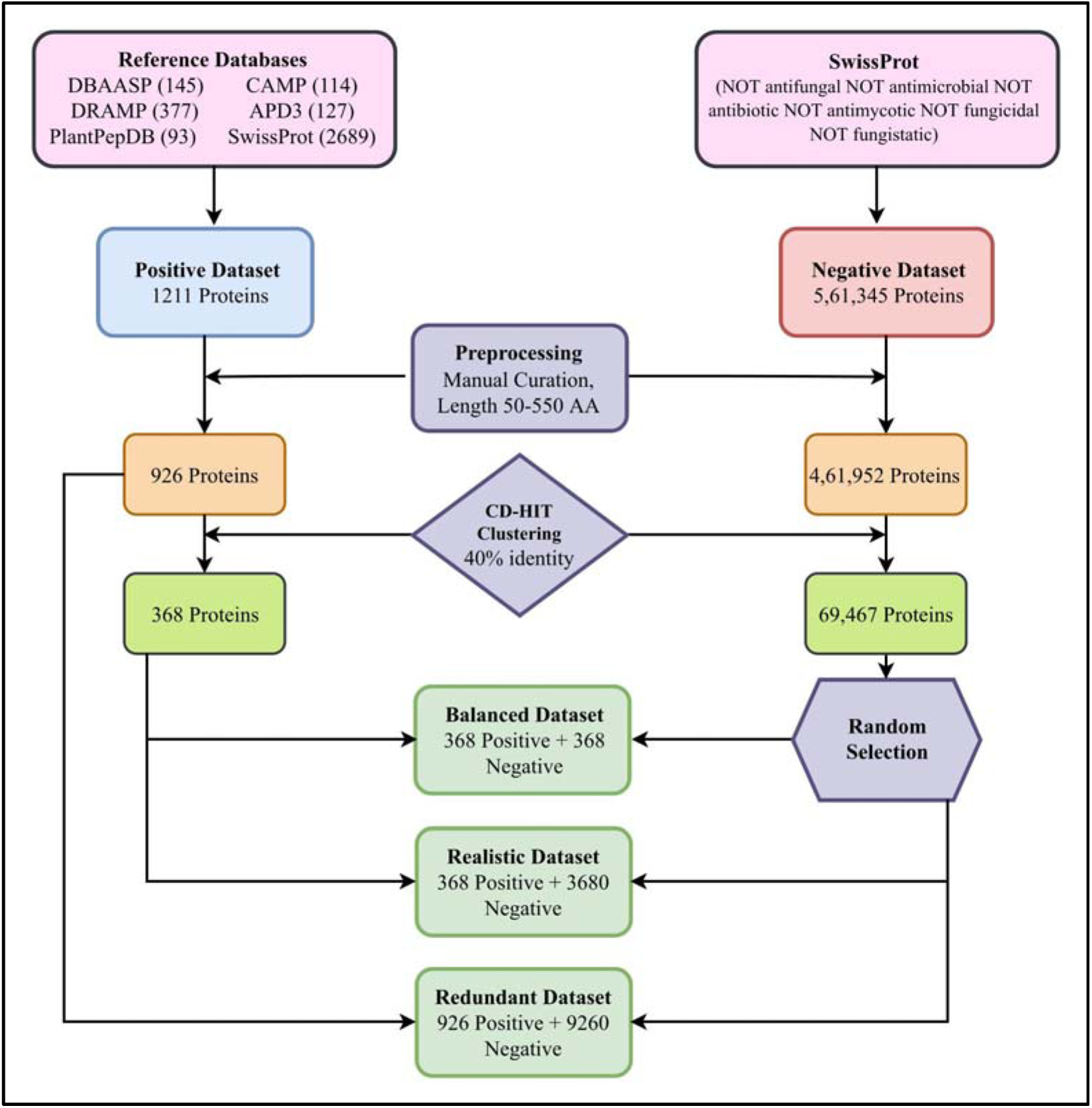
Diagrammatic representation of dataset creation.

All datasets were divided into an 80:20 ratio, where 20% of the sequences were reserved as an independent validation set. The remaining 80% constituted the training set and was used for five-fold cross-validation, model development, and hyperparameter optimisation.

### 5.2 BLAST for Similarity Search

The Basic Local Alignment Search Tool (BLAST) is heavily used for the annotation of protein and nucleotide sequences based on sequence similarity. Initially, we created a local BLAST reference database based on the training dataset using the *makeblastdb* program. Then, we have attempted to identify antifungal proteins and non-antifungal proteins using BLASTP, which queries protein sequences from the validation database against the local BLAST database to find similarity and assign annotation based on the matches. The BLASTP was run on various E-value thresholds from 10 to 0.00001 to decide the best threshold based on prediction accuracy.

The antifungal and non-antifungal were predicted based on the single best hit obtained for each query sequence; the reference database had a suffix _1 for positive proteins and _0 suffix for negative proteins after the sequence ID for convention.

### 5.3 Motif analysis

The motif discovery analysis was performed on antifungal proteins by using Motif-EmeRging and with Classes-Identification (MERCI) tool [27], software to find motifs in protein sequences. Motif analysis provides information regarding recurring sequence patterns found in the provided antifungal protein sequences. We used two classification systems: the first is Koolman-Rohm, and the second is None. In combination with various numbers of motifs (-k) and minimum frequency for positive sequences (-fp) thresholds, we set optimised parameters to find the optimal parameters. The best parameters were selected based on % accuracy.

### 5.4 Composition-based features

The standalone Pfeature [28] tool was used to generate 15 different types of compositional features, totalling 959 features. We also calculated the same features after splitting the sequence into four equal parts, generating 3836 features. The detailed features are mentioned in **Supplementary Table 1.**

### 5.5 Probability-based features

To generate probability-based features from the original Amino Acid Composition (AAC) and Pseudo Amino Acid Composition (PAAC) feature sets, a feature-wise transformation was performed using logistic regression. For each individual feature column in the training data, a distinct logistic regression model was trained to predict the class label based solely on that single feature. Subsequently, this trained model was used to calculate the class probability for each instance in both the training and validation sets. This process was repeated for every feature, creating new datasets where each original feature value was replaced by its corresponding probability score, effectively normalising the feature space and encoding the predictive power of each component into a probabilistic value between 0 and 1.

### 5.6 Evolutionary-based features

The evolutionary features were extracted from the protein sequences to try and capture the conserved information in the sequences. This was accomplished by using a position-specific scoring matrix (PSSM) algorithm provided by BLAST variant Position-Specific Integrated BLAST (PSI-BLAST) [29]. The PSI-BLAST was used to generate a profile of 20X20 for each sequence, using SWISSPROT as the reference database [24]. These profiles were then passed to the POSSUM package, and the ‘*pssm_composition*’ module was used to generate 400 evolutionary-based features [30]. These features were then used to train machine learning models.

### 5.7 Feature selection

Using high-dimensional feature vectors is not always the best choice in machine learning, as they will lead to overfitting, unnecessary calculations, and increased prediction time. There are multiple methods available for feature selection; we implemented the ones known to work well for biological data, such as the forward sequential feature selection (FSFS) technique, which starts with an empty feature set, then tests all single features, and selects the best one. It keeps adding features one by one till the performance no longer improves. Support vector classifier (SVC)-L1-based feature selection technique, and minimum redundancy maximum relevance (MRMR) technique were used with different minimum feature selection thresholds, and models were trained on them to see the results.

### 5.8 Machine learning based classification

We applied various machine learning techniques to classify between antifungal and non-antifungal proteins. The logistic regression (LR), Support Vector Machine (SVM), K-nearest neighbour (KNN), Random forest (RF), Gradient boosting (GB), Adaptive Boosting (AdaB), Bernoulli Naive Bayes (BNB), Gaussian Naive Bayes (GNB), Decision Tree (DT), eXtreme Gradient Boosting (XGB) and Extra tree (ET) classifiers were used to develop a classification model using default parameters (random_state = 1) as a preliminary phase. Later, the best five models based on their performance in the preliminary were selected, and a grid search algorithm was used for hyperparameter optimisation.

### 5.9 Large Language-based model

Large language models were also implemented to improve the classification of antifungal proteins. We used multiple pretrained models, such as Evolutionary Scale Modelling (ESM2) [31] variants t6_8M, t12_35M, t33_650M, and t36_3B, and ProtBERT [32]. Each model was further fine-tuned using embeddings extracted from antifungal and non-antifungal proteins for better prediction accuracy. The embeddings were also used as features for training the machine learning models.

### 5.10 Ensemble model

To improve the accuracy of the predictions even further, we used ensemble modelling, which is based on a weighted-scoring approach. We integrated three different methods, namely 1) Similarity-based method, 2) Motif-based method and 3) Machine learning–based or Large Language Model–based method. This method first applies BLAST on the given query protein sequence (from the validation dataset) and finds its best hit under an optimised e-value. If the sequence matches with an antifungal protein, we add +0.5 to the prediction probability generated by the ML/LLM model; if it matches with a non-antifungal protein, we subtract 0.5 from the prediction probability. Secondly, we use MERCI to implement motif search. If known antifungal motifs are found, we again add +0.5 to the probability. Finally, we apply the machine learning–based or large language model–based method to generate the base probability, and then sum the contributions from the previous methods to decide the final prediction

### 5.11 Cross-validation

In this study, each of the three data sets was divided into an 80:20 ratio, using 80% for training and 20% for validation. In the training set, we employed a five fold cross validation (CV) protocol: the training dataset was divided into five distinct folds, and four folds were used iteratively to fit models and test on the remaining fold, with each fold acting as the validation subset once. In particular, we evaluated our models based on the area under the receiver operating characteristic curve (AUC-ROC) as a threshold-independent measure, and the Matthews correlation coefficient (MCC) as a threshold-dependent measure. The equations for these calculations are mentioned in previous studies [20]. Additionally, other popular performance metrics like sensitivity, specificity and accuracy were also calculated to evaluate the models. The diagrammatic representation of the methodology used to develop AntiFP2 is provided in **Figure 4**.

**Figure 4:**
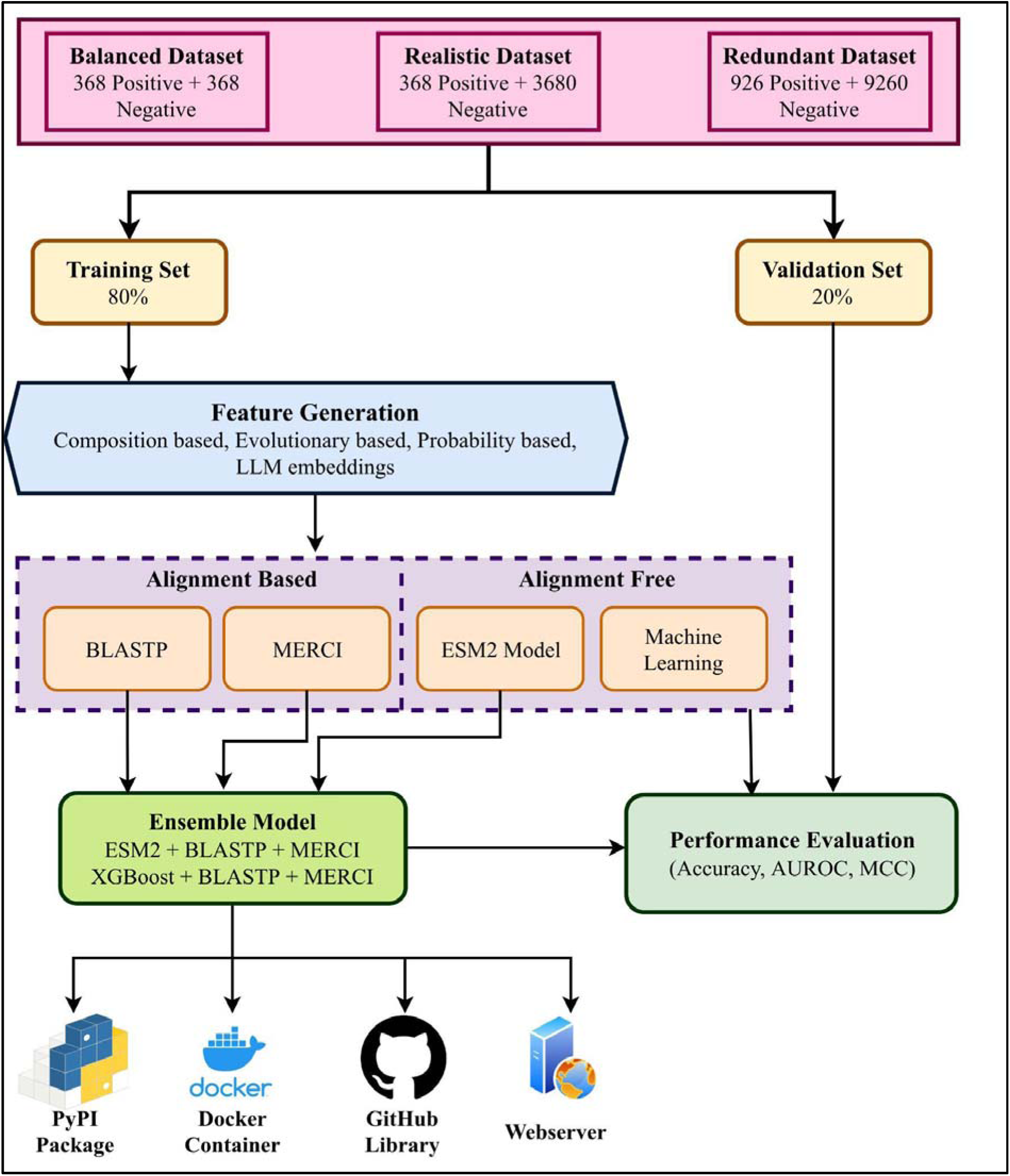
Diagrammatic representation of the architecture of AntiFP2.

### 5.12 Webserver Architecture

In this study, we also developed a webserver, AntiFP2, for antifungal protein prediction (https://webs.iiitd.edu.in/raghava/antifp2/). The platform is implemented using HTML5, CSS, Java, and PHP, providing a responsive and device-independent interface accessible from desktops, tablets, and mobile devices. The server features a Predict module that enables users to submit protein sequences and obtain antifungal protein predictions using the AntiFP2 models in a user-friendly manner.

### 5.13 Genome-wide and metagenome antifungal prediction module

To enable antifungal protein identification directly from genomes and metagenomes, we developed an automated prediction module that integrates PROKKA v1.14.5 [33] and SeqKit v2.10.0 [34] with the AntiFP2 model. The workflow accepts assembled contigs, performs gene prediction and protein annotation using PROKKA [33], classifies the resulting proteins with AntiFP2, and uses SeqKit to extract sequences predicted as antifungal. This module provides a streamlined, reproducible pipeline for large-scale genome- and metagenome-wide AFP identification with minimal user input.

## Supporting information

Supplementary_Antifp2.xlsx

## Ethics approval and consent to participate

Not applicable.

## Consent for publication

Not applicable.

## Availability of data and materials

All the datasets used in this study are available at the “AntiFP2” web server, https://webs.iiitd.edu.in/raghava/antifp2/dataset.html. A metagenome and genome-level annotation pipeline is provided as a standalone workflow on GitHub (https://github.com/raghavagps/antifp2) and as an environment-independent Docker container (https://hub.docker.com/r/pratik0297/antifp2), offering an easy installation option for all users. The balanced, realistic and redundant datasets are available on (https://webs.iiitd.edu.in/raghava/antifp2/dataset.html). The fine-tuned ESM2 model is available on Hugging Face (https://huggingface.co/raghavagps-group/antifp2).

## Authors’ contributions

PS, RT collected and processed the datasets. PS, SC implemented the algorithms and developed the prediction models. PS, SC, GPSR analyzed the results. SC created the front-end and back-end of the web server. PS, SC, GPSR penned the manuscript. GPSR conceived and coordinated the project. All authors have read and approved the final manuscript.

## Acknowledgments

Authors are thankful to the Indraprastha Institute of Information Technology, Delhi and Department of Science and Technology (DST-INSPIRE) for fellowships and financial support, and the Department of Computational Biology, IIITD, New Delh,i for infrastructure and facilities. We would like to acknowledge that the Figures were created using Draw.io.

